# Tau-Tubulin Kinase 2 restrains microtubule-depolymerizer KIF2A to support primary cilia growth

**DOI:** 10.1101/2023.02.21.529351

**Authors:** David Vysloužil, Ondřej Bernatík, Tereza Renzová, Lucia Binó, Andrea Lacigová, Lukáš Čajánek

**Affiliations:** Laboratory of Cilia and Centrosome Biology, Department of Histology and Embryology, Faculty of Medicine, Masaryk University, Kamenice 3, 62500 Brno, Czech Republic; Department of Experimental Biology, Faculty of Science, Masaryk University, Kamenice 5, 62500 Brno, Czech Republic

## Abstract

The initiation of assembly of primary cilia, organelles with crucial functions in development and disease, is under the control of Tau-Tubulin Kinase 2 (TTBK2). Recent work has implicated TTBK2 also in the regulation of primary cilia maintenance and function. However, the mechanisms underlying individual functions of TTBK2 in primary cilia are not fully understood.

Here, to dissect the role of TTBK2 in primary cilia maintenance in human cells, we examined disease related TTBK2 truncations. We demonstrate that these truncated protein moieties show selective activity towards TTBK2 substrates. This creates a semi-permissive condition where partial TTBK2 activity suffices to support the initiation of ciliogenesis but fails to sustain primary cilia length. Subsequently, we show that the defects in primary cilia growth are linked to aberrant turnover of kinesin KIF2A at basal body. Furthermore, we demonstrate that TTBK2 regulates KIF2A by phosphorylation, which in turn restrains its levels at the ciliary base to promote primary cilia elongation and maintenance.

Taken together, our data highlight the regulation of KIF2A by TTBK2 as an important mechanism governing primary cilia in human cells.

## Introduction

Primary cilia are evolutionarily conserved signalling organelles found on the surface of a broad spectrum of eukaryotic cells^1,2^. They play major role in intercepting extracellular stimuli, which helps to govern complex processes in multicellular organisms, including humans^3,4^. Even though these organelles have not attracted much attention in the past, their biomedical importance became quite apparent following the discovery of ciliopathies, an expanding group of human diseases caused by ciliary dysfunction^5,6^.

Primary cilium typically appears as a hair-like protrusion of the cell membrane. It consists of the mother centriole (MC)-derived basal body, the transition zone allowing sorting of ciliary components, and the microtubule-based axoneme enclosed within a ciliary membrane^7^. The MC distal end is decorated by two sets of appendage proteins. The distal set of appendages is especially important for the initiation of cilia formation – it helps to dock the MC to preciliary vesicles and/or cell membrane and recruits pro-ciliogenesis factors to the base of the growing cilium^8,9^. To initiate the outgrowth of ciliary axoneme, capping complexes consisting of CP110 and CEP97 proteins are removed from the MC distal end^10^. Subsequently, the axoneme extends with the assistance of intraflagellar transport (IFT) machinery^11^.

Perhaps the most prominent regulator of the cilium assembly pathway outlined above is Tau-Tubulin Kinase 2 (TTBK2)^12^, a serine/threonine kinase from the CK1 superfamily^13,14^. To initiate ciliogenesis, TTBK2 is recruited to the distal appendages via interaction with CEP164^25^ Thus, TTBK2 gets in prime position to govern key events at the base of the future cilium by its kinase activity^12^. The processes regulated by TTBK2 include the recruitment of Golgi-derived preciliary vesicles^15^ and IFT proteins^12^, or the removal of CP110/CEP97^12^.

However, the full scope of TTBK2-regulated mechanisms and their relation to ciliogenesis remains to be discovered. TTBK2 has been shown to phosphorylate a number of MC-related (e.g. CEP164, CEP83, CEP89^15,16^) or cell signalling-related proteins (e.g. Dishevelled, a component of the WNT signaling pathway^16,17^), but the functional relevance of that is often not clear. Furthermore, recent work highlights the possibility of TTBK2 maintaining length and integrity of primary cilia^18,19^, but molecular mechanisms that would link TTBK2 to axoneme length control remain elusive. Finally, truncating mutations in TTBK2 have been shown to trigger the onset of spinocerebellar ataxia 11 (SCA11), a rare progressive neurodegenerative disease^19–21^. While the genetic link between TTBK2 and SCA11 has been well described, the molecular pathology of SCA11 is unclear.

In this study, we aimed to elucidate the role TTBK2 plays beyond initiating cilia formation in human cells. As targeting TTBK2 through conventional loss-of-function approaches completely blocks cilia formation^12,16^, we examined TTBK2 variants, similar in length and sequence to the SCA11 mutants^20^. We show that expressing truncated TTBK2 in TTBK2 null hTert-RPE-1 cells creates unique semi-permissive conditions for ciliogenesis. In turn, we utilize this model system to reveal a TTBK2-mediated regulatory mechanism involving kinesin KIF2A, a member of kinesin-13 family with microtubule-depolymerizing activity^22,23^. KIF2A has been previously shown to aid the resorption of primary cilia following cell cycle re-entry^24^, but its role during cilia assembly was not clear. Here, we demonstrate that the interplay between KIF2A and TTBK2 represents an important regulatory mechanism governing primary cilia formation in human cells.

## Results

### Truncated TTBK2 exhibits reduced activity

To establish a system with reduced TTBK2 activity, we focused on TTBK2 truncations. The identified *TTBK2* frameshift mutations lead to SCA11 pathology in human^20^. In mice, the corresponding truncated protein moieties showed significantly reduced or no activity^12,19^. First, we examined the biochemical activity of TTBK2 construct truncated shortly after the kinase domain (1-450 AA, referred to as TTBK2^trunc1^). To this end, we co-expressed Myc-tagged N-terminal part of CEP164 (1-467 AA, CEP164^NT^) with Flag-TTBK2^wt^ or Flag-TTBK2^trunc1^ in wild-type HEK293T cells. Consistently with the reported TTBK2 phosphorylation of CEP164^16,25^, we observed that Flag-TTBK2^wt^ induced a mobility shift in MYC-CEP164^NT^(1A). In contrast, the expression of Flag-TTBK2^trunc1^ showed no effect. Next, we examined the phosphorylation of S201 in CEP164, a site targeted by TTBK2^16^. In comparison to wild-type hTERT RPE-1 cells (S1A-B), we found MC levels of phosphorylated S201 (pCEP164) reduced in CEP164 KO and TTBK2 KO, respectively, as expected. Having confirmed the specificity of the pCEP164 antibody, we in turn examined pCEP164 levels at MC in hTERT RPE-1 TTBK2 KO cells DOX-inducibly expressing individual Flag-tagged variants of TTBK2. Using this system, we found pCEP164 MC signal markedly increased in cells expressing Flag-TTBK2^wt^, compared to Flag-TTBK2^kd^ or Flag-TTBK2^trunc1^-expressing cells (1B-C), in line with our WB data.

To corroborate our findings, we next examined phosphorylation of Dishevelled-3 (DVL3). As expected, DVL3 was phosphorylated when co-expressed with Flag-TTBK2^wt^, but not Flag-TTBK2^kd^, as indicated by the observed DVL3 mobility shift in HEK293T cells (1D). Intriguingly, we also observed mobility shift of DVL3 when co-expressed with Flag-TTBK2^trunc1^ (1D), or GFP-TTBK2^trunc2^ (S1C, an insertion of adenosine at nucleotide 1329 leads to premature stop codon formation^20^ and in turn expression of a 1-450 AA long protein). In line with existing data^21^, we noted that TTBK2^trunc1/2^ protein products were more abundant than TTBK2_wt_ (1A+D). We titrated TTBK2 plasmids and confirmed that both TTBK2_wt_ and TTBK2_trunc1_ induce DVL3 mobility shift also when expressed at comparable levels (S1D). To support our findings further, we examined the subcellular localization of overexpressed DVL3. Depending on its phosphorylation status, DVL3 can form dynamic protein assemblies^17,26–28^. Indeed, while we typically found DVL3 localized in cytosolic punctae in TTBK2 KO, it showed even distribution when Flag-TTBK2^wt^ or Flag-TTBK2^trunc1^ were expressed (1E). Further, we examined the activity of truncated TTBK2 using a dual luciferase WNT pathway reporter assay TOPFLASH^29,30^. Interestingly, expression of beta-catenin (BCAT) with a S37A activating mutation^31^ in HEK293T led to strong activation of the WNT reporter, which was attenuated by both Flag-TTBK2^wt^ and Flag-TTBK2^trunc1^, but not by Flag-TTBK2^kd^ (1F).

In sum, our results suggested that the tested truncated TTBK2 moieties possess biological activity towards a subset of TTBK2 substrates rather than being completely inactive.

### Truncated TTBK2 is sufficient to trigger cilia assembly

To test the ability of truncated TTBK2 to affect ciliogenesis, we first examined its effect on the removal of CP110 from the MC and the recruitment of IFT88 to the base of the cilium. Interestingly, we found both Flag-TTBK2^wt^ and Flag-TTBK2^trunc1^, but not Flag-TTBK2^kd^, to efficiently induce the MC removal of CP110 in TTBK2 KO (2A-B). In addition, while Flag-TTBK2^kd^ failed to promote the IFT88 MC recruitment, Flag-TTBK2^wt^ and to a lesser extent also Flag-TTBK2^trunc1^ showed a rescue effect on IFT88 MC levels (2C-D), supporting the notion that TTBK2^trunc1^ was partially active.

In turn, we tested the ability of TTBK ^trunc1/2^ to support primary cilia formation. First, we confirmed that while cells lacking TTBK2 were completely devoid of primary cilia positive for ARL13B^32^, Flag-TTBK2^wt^ but not Flag-TTBK2^kd^ rescued ciliogenesis (2E, S2A). Remarkably, TTBK2 KO cells expressing either Flag-TTBK2^trunc1^ or Flag-TTBK2^trunc2^ formed significantly shorter primary cilia (2E-F) and showed lower or similar efficiency of primary cilia formation (S2A). In support of that, we observed a similar rescue effect on ciliogenesis for Flag-TTBK2^wt^ and Flag-TTBK2^trunc1^ also in HEK293T TTBK2 KO cells (S2B). Furthermore, a chimeric construct consisting of TTBK2^trunc1^ and the C-terminal part of CEP164 (468-1460 AA), which we termed CEP164 chimera, rescued both cilia number and cilia length to the level of TTBK2^wt^ (2E-F, S2A). This data demonstrated that the observed defects in primary cilia assembly can be effectively rescued by re-targeting TTBK2^trunc1^ to the basal body.

### TTBK2 truncation disrupts axoneme growth

Given the rather modest defect in IFT88 MC levels in TTBK2^trunc1^-expressing cells, we hypothesized that TTBK2 might regulate axoneme length by an additional mechanism, distinct from its role in IFT recruitment. To this end, we employed expansion microscopy to examine the ultrastructure of primary cilia induced by TTBK2^wt^ vs TTBK2^trunc1/2^. Primary cilia formed in the TTBK2^wt^ condition showed a typical arrangement of acetylated tubulin (AcTUB) positive axonemal microtubules, enclosed in a ciliary membrane labelled by ARL13B (3A). Many cilia found in the TTBK2^trunc1^ condition were similar to that, just with a shorter axoneme. However, we also often noticed structures consisting of ARL13B-positive vesicles docked to the MC distal end, with no apparent sign of axonemal microtubules extension (3A, panel on the right).

To assess the extent of this defect, we determined a ratio of combined MC + axoneme length relative to the length of the daughter centriole. For centriole pairs with no ARL13B signal, the length ratio was around 1. Given that, we considered a value between 1 to 1.5 as indication that no axoneme formed. As expected, this ratio markedly shifted in ARL13B-positive centriole pairs in TTBK2^wt^, as one centriole (presumably the mother) templated the axoneme. While we found very few ARL13-positive centriole pairs here with a ratio between 1 and 1.5 (3 out of 19), the corresponding population of ARL13B-positive centrioles with no apparent axoneme extension was more abundant (14 out of 45) in TTBK2^trunc1^ (3B). In addition, the determined ratio was typically smaller in TTBK2^trunc1^, in agreement with the observed cilia length defect. In sum, our data indicated that the activity of truncated TTBK2 is sufficient to trigger early events of cilia formation but fails to sufficiently support later stages.

Given the phenotypes we found, we next examined the dynamics of primary cilia formation in TTBK2^wt^ and TTBK2^trunc1^, respectively, using live-cell imaging microscopy. TTBK2 KO hTERT RPE-1 expressing Flag-TTBK2^wt^ formed steadily growing NeonGreen-tagged ARL13B cilia during the 5h period of the experiment (3C-D). In contrast, cilia detected in Flag-TTBK2^trunc1^-expressing TTBK2 KO cells elongated very slowly, sometimes even shortened towards the end of the experiment. We also noted a modest increase in cilia breakage events (3E) in the TTBK2^trunc1^ condition (0.355 breaks per cilium) compared to TTBK2^wt^ (0.222 breaks per cilium).

Taken together, our data revealed a prominent axoneme elongation defect related to aberrant activity of truncated TTBK2. Thus, we concluded that TTBK2 might govern axoneme growth by regulating the stability of its microtubules.

### KIF2A overactivation phenocopies TTBK2 truncation

Having linked TTBK2 activity to the dynamics of primary cilia, we sought to reveal the underlying mechanism. We focused on KIF2A, a kinesin with microtubule-depolymerizing activity, previously implicated in cilia resorption upon cell cycle re-entry^24^ and phosphorylated by TTBK2^22^.

First, we generated hTERT RPE-1 cell lines DOX-inducibly expressing GFP-KIF2A^wt^ or GFP-KIF2A^KVD^ (a mutant defective in depolymerizing microtubules^24,33^). We found both KIF2A constructs to preferentially localize to centrioles and their proximity (4A). Intriguingly, we could occasionally detect a signal of GFP-KIF2A^KVD^ also decorating the axonemes of primary cilia. In addition, we resolved the signal of GFP-KIF2A^wt^ using expansion microscopy and found it to form ring-like structures near the MC distal end, closely resembling the localization pattern of distal or subdistal appendage proteins (4B). Furthermore, our expansion microscopy protocol also revealed a faint signal of GFP-KIF2A^wt^ decorating the AcTUB+ ciliary axoneme. Noteworthy, we also detected endogenous KIF2A localized to centrioles and their proximity (4C-D), which was abolished by paclitaxel treatment (5µM, 4h), suggesting a dependency on intact microtubules.

In turn, we examined the effects of KIF2A on primary cilia formation. Following DOX-induced expression of GFP-KIF2A^wt^, we could readily observe a reduction in cilia length (4E-F), in line with previous work^24^. In addition, we found that GFP-KIF2A^wt^, but not GFP-KIF2A^KVD^, reduced the percentage of ciliated cells (4G).

Having established that elevated KIF2A levels phenocopy the cilia defects related to TTBK2^trunc1/2^, we examined a possible causality between KIF2A and TTBK2^trunc1/2^. Intriguingly, we found that endogenous KIF2A MC levels were markedly elevated in TTBK2 KO cells expressing Flag-TTBK2^trunc1^ or Flag-TTBK2^trunc2^ compared to Flag-TTBK2^wt^ or Flag-TTBK2^kd^ (4H-I). We confirmed this was specific to DOX-induction of FLAG-TTBK2^trunc1^ and not due to a simple cell line-to-cell line variability (S3A-B). Next, following the validation of KIF2A siRNA efficacy (S3C), we in turn tested if KIF2A depletion rescues any of the cilia defects associated with truncated TTBK2. Remarkably, even though KIF2A depletion did not significantly change the percentage of ciliated cells (S3D), it fully rescued the cilia length defect in Flag-TTBK2^trunc1^-expressing cells (4J-K). Importantly, KIF2A depletion showed no additive effect on primary cilia length in the TTBK2^wt^ condition.

To conclude, our data established that the activity of KIF2A in primary cilia interferes with cilia assembly and in turn mediates some of the defects related to TTBK2^trunc1/2^.

### TTBK2-induced phosphorylations alter KIF2A basal body pool

To reveal how TTBK2 regulates KIF2A, we first examined KIF2A phosphorylation induced by TTBK2. Following co-expression of Flag-KIF2A^wt^ with GFP-tagged TTBK2 variants in HEK293T TTBK2 KO we immunopurified Flag-KIF2A^wt^ and subjected the isolates to MS/MS analysis. With this approach, we identified several S/T phosphosites induced by GFP-TTBK2^wt^, but not GFP-TTBK2^trunc2^ or GFP-TTBK2^kd^ (5A). We focused on S137 and S140, as they were in proximity to the previously reported phosphosite S135^22^, and were evolutionary conserved. As we observed TTBK2^trunc1/2^-mediated increase in the levels of KIF2A at the MC and its proximity, but no difference in total levels of KIF2A (S3A), we reasoned TTBK2 might regulate KIF2A recruitment. To examine a possible function of S135, S137, and S140 phosphorylation in KIF2A recruitment, we expressed GFP-KIF2A^wt^, GFP-KIF2A^PhoMim^ (S135, S137, and S140 mutated to Q) or GFP-KIF2A^PhoDead^ (S135, S137, and S140 mutated to A) in hTERT RPE-1. Intriguingly, we found GFP-KIF2A^wt^ and GFP-KIF2A^PhoDead^, but not GFP-KIF2A^PhoMim^, readily localizing to the MC and its proximity (5B-C). This suggested that TTBK2 phosphorylation of S135-S140 likely facilitates ciliogenesis by preventing the accumulation of KIF2A at MCs. However, our results also hinted at possible limitations of our experimental setup, as we were not able to resolve a difference in MC levels between GFP-KIF2A^wt^ and GFP-KIF2A^PhoDead^. To support the role of TTBK2 phosphorylation in KIF2A turnover by other means, we carried out FRAP analysis of GFP-KIF2A^wt^. Importantly, when comparing the effect of Flag-TTBK2^wt^ and Flag-TTBK2^trunc1^ in TTBK2 KO, we found the recovery halftime (Thalf) of GFP-KIF2A^wt^ at the MC significantly slower in TTBK2^trunc1^ condition (5D-E).

Taken together, our results suggest that TTBK2 phosphorylation destabilizes KIF2A basal body pool and prevents its accumulation at the base of the cilium, which supports efficient axoneme growth. In contrast, truncated TTBK2 inefficiently phosphorylates KIF2A, which seems to alter its turnover, leading to increased KIF2A levels at the ciliary base, and in turn to reduced ciliary length (Figure 6).

**Fig 1:**
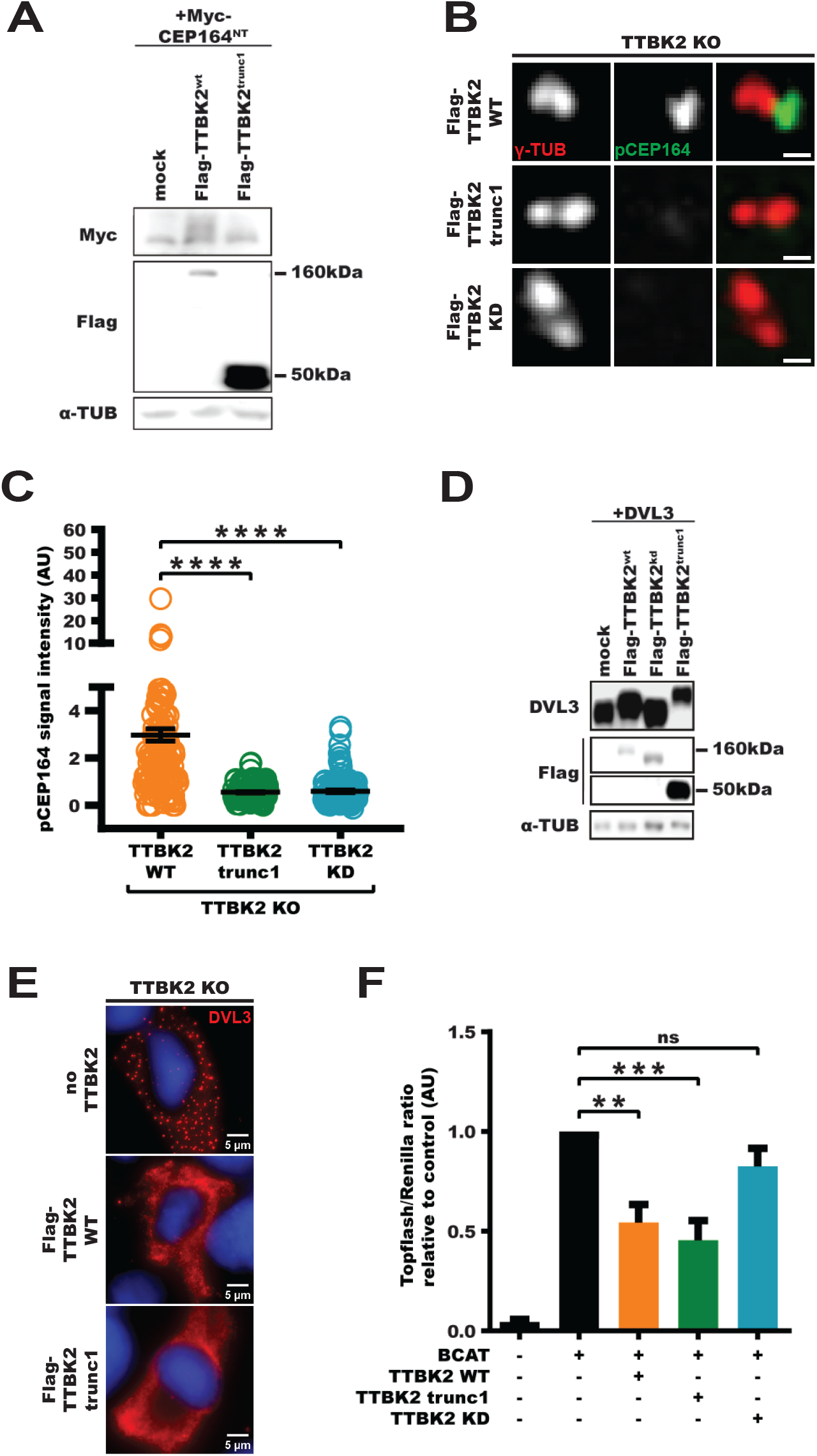
Truncated TTBK2 exhibits reduced activity. Western blot analysis of lysates from HEK293T cells transfected with Myc-CEP164^NT^ and Flag-TTBK2 constructs. (B,C) hTERT RPE-1 cells stably expressing Flag-TTBK2 constructs were fixed and stained for pCEP164, scale bars: 0.5µm. Relative centriolar levels of pCEP164 (normalized to γ-TUB, one-way ANOVA, \*\*\*\**P*<0.0001) are quantified in C. (D) Western blot analysis of lysates from HEK293T cells transfected with DVL3 and Flag-TTBK2 constructs. (E) hTERT RPE-1 cells expressing Flag-TTBK2 constructs were transfected with DVL3, fixed, and stained for DVL3. (F) Topflash dual-luciferase assay in HEK293T cells transfected with the indicated plasmids. Topflash reporter signal was normalized to that of Renilla reporter, the obtained values were related to BCAT condition (which was set to 1), one-way ANOVA, \*\**P*<0.01, \*\*\**P*<0.001).

**Fig 2:**
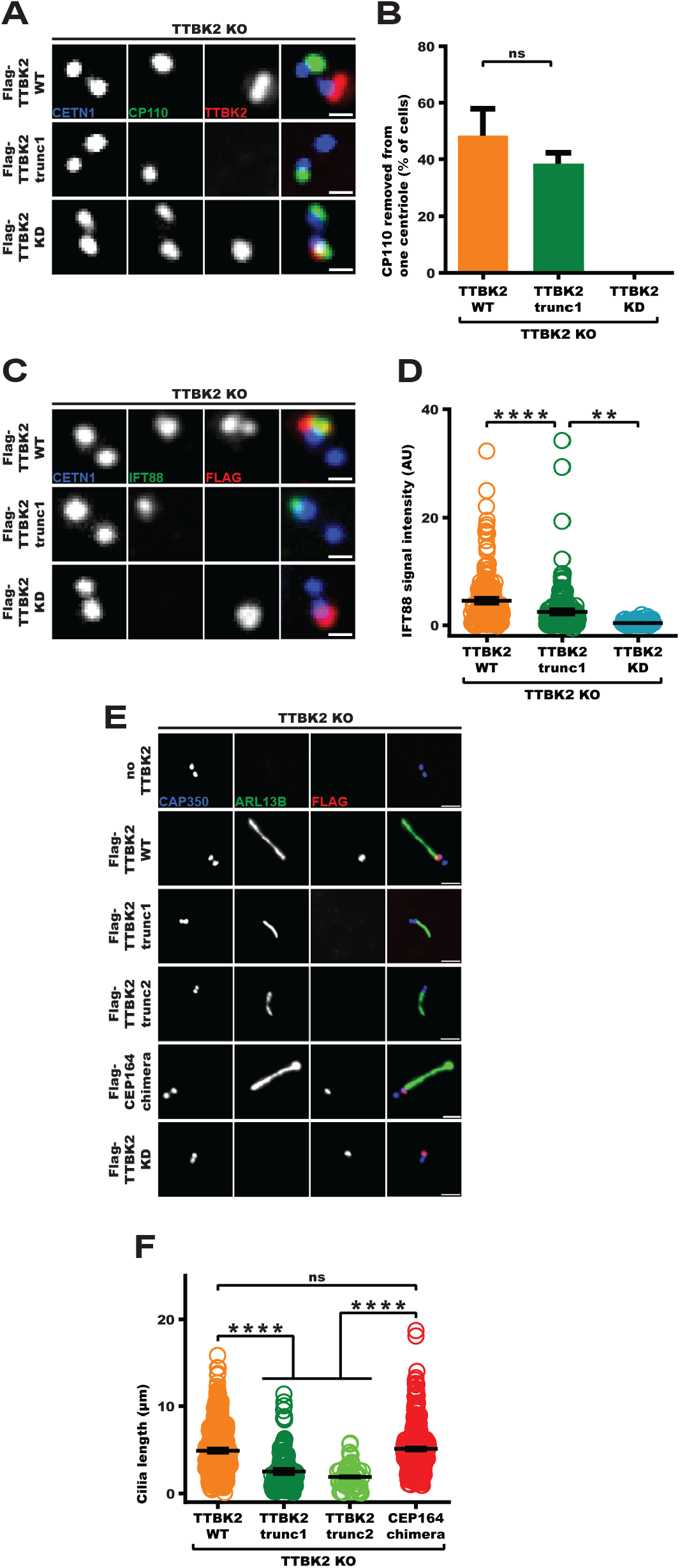
Truncated TTBK2 is sufficient to trigger cilia assembly. (A,B) hTERT RPE-1 cells expressing Flag-TTBK2 constructs were fixed and stained for CP110 and the indicated antibodies (scale bars: 0.5µm). Centriole pairs with one vs two CP110 signals (unpaired T-test, *P*=0.2153) are quantified in B. (C,D) hTERT RPE-1 cells expressing Flag-TTBK2 constructs were fixed and stained for IFT88 and the indicated antibodies (scale bars: 0.5µm). Relative centriolar levels of IFT88 (normalized to CETN1, one-way ANOVA, \*\*\*\**P*<0.0001, \*\**P*<0.01) are quantified in D. (E,F) hTERT RPE-1 cells expressing TTBK2 constructs were fixed and stained for ARL13B and the indicated antibodies (scale bars: 2µm). The length of cilia (one-way ANOVA,\*\*\*\**P*<0.0001) is quantified in F.

**Fig 3:**
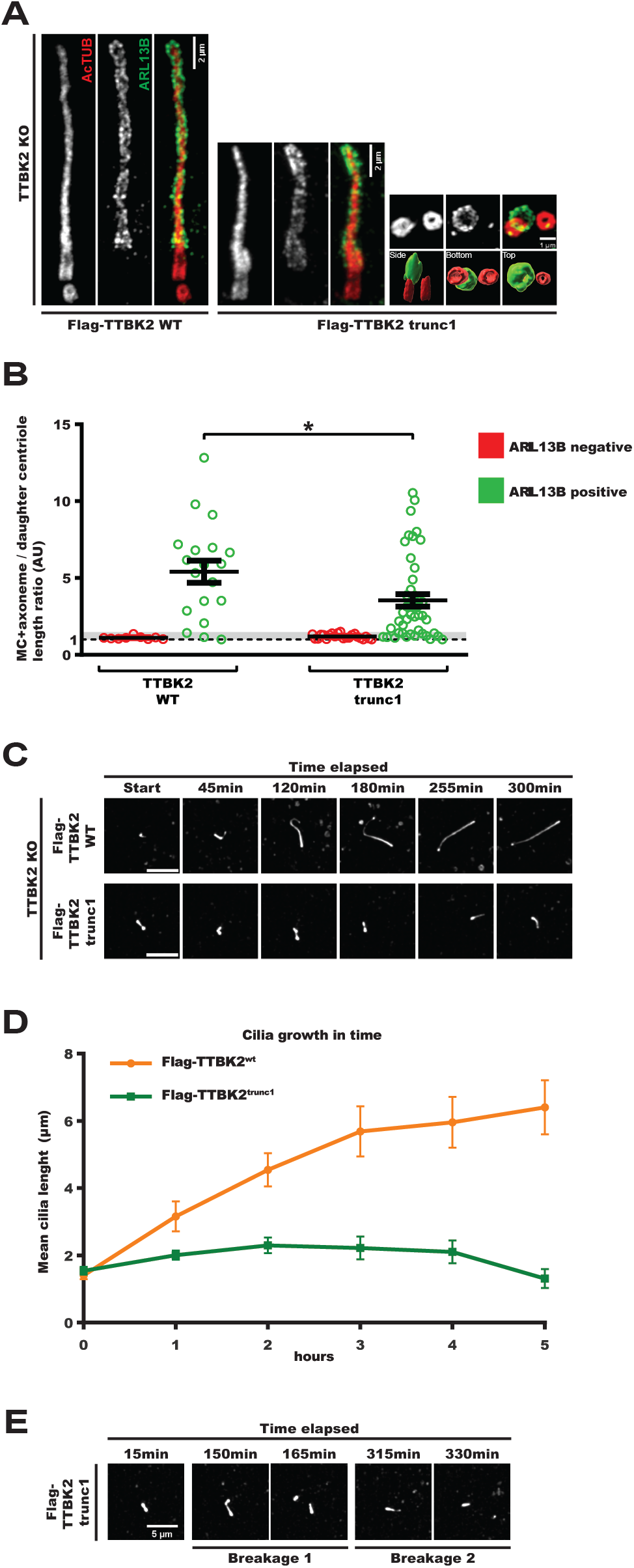
TTBK2 truncation disrupts axoneme growth. (A,B) hTERT RPE-1 cells expressing Flag-TTBK2 constructs were subjected to expansion microscopy and stained for AcTUB and ARL13B. Bottom right panel shows Imaris 3D reconstructions viewed from different perspectives. The length ratios of MC+axoneme to DC (unpaired T-test, \**P*<0.05) are quantified in B, values between 1 and 1.5 are indicated by the grey stripe. (C,D) hTERT RPE-1 cells expressing NeonGreen-ARL13B and Flag-TTBK2 constructs were subjected to live-cell imaging (images show the NeonGreen signal, scale bars: 5µm). The mean length of cilia measured over the coarse of 5 hours is quantified in D. (E) Cells expressing NeonGreen-ARL13B and Flag-TTBK2^trunc1^ exhibited frequent cilia breakage events.

**Fig 4:**
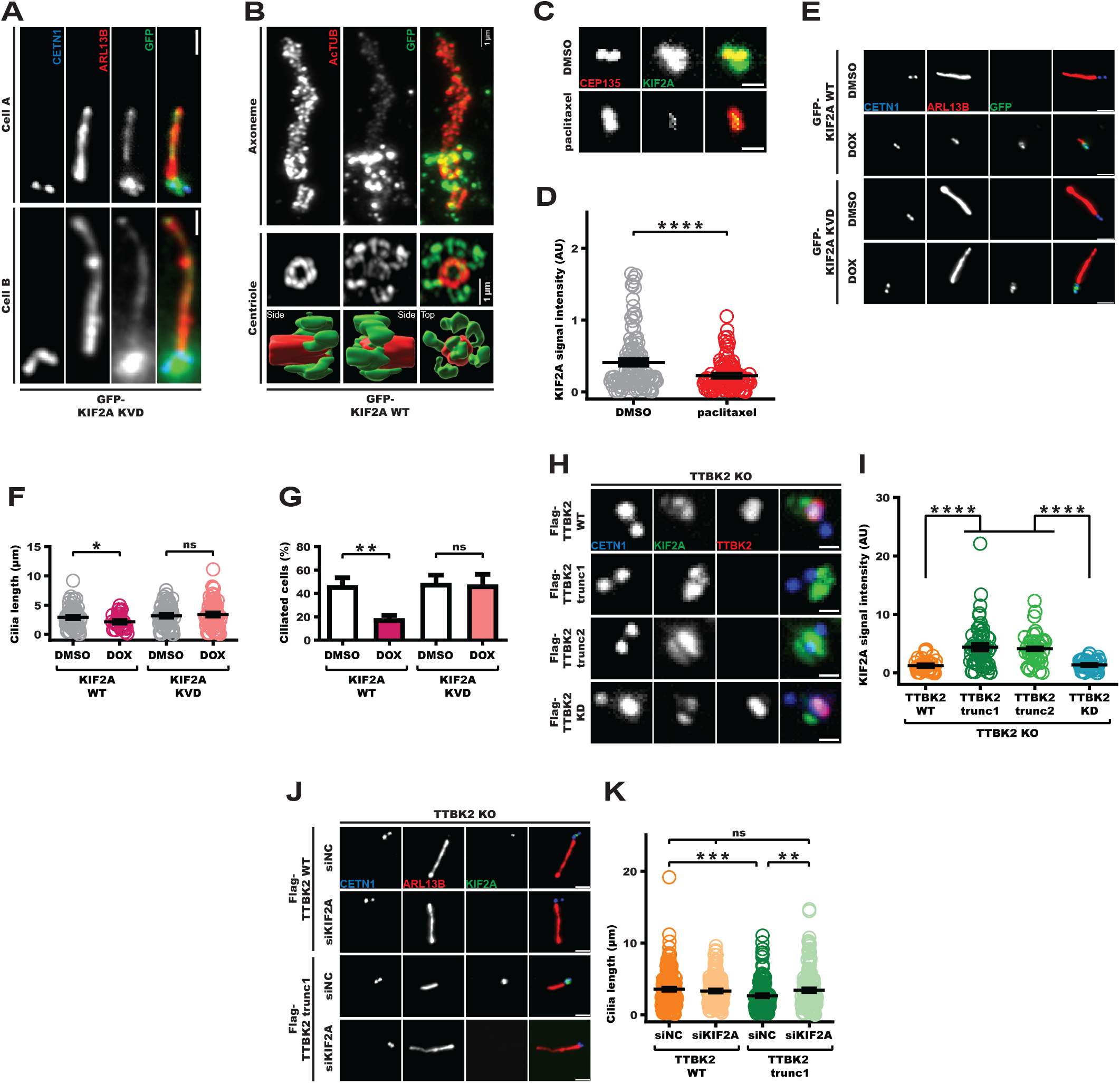
KIF2A overactivation phenocopies TTBK2 truncation. (A) hTERT RPE-1 cells expressing GFP-KIF2AKVD were fixed and stained for ARL13B and CETN1 (scale bars: 1µm). (B) hTERT RPE-1 cells expressing GFP-KIF2A^wt^ were fixed, subjected to expansion microscopy, and stained for AcTUB and GFP. Bottom panel shows Imaris 3D reconstructions viewed from different perspectives. (C,D) hTERT RPE-1 cells were treated with DMSO (control group) or paclitaxel (5µM, 4h) and stained for KIF2A and the indicated antibodies (scale bars: 1µm). Relative centriolar levels of KIF2A (normalized to CEP135, unpaired T-test, \*\*\*\**P*<0.0001) are quantified in D. (E-G) hTERT RPE-1 cells expressing GFP-KIF2A constructs were treated with DMSO or DOX, fixed, and stained for ARL13B and the indicated antibodies (scale bars: 2µm). Cilia length and number (unpaired T-tests, \**P*<0.05, \*\**P*<0.01) are quantified in F and G respectively. (H,I) hTERT RPE-1 cells expressing Flag-TTBK2 constructs were fixed and stained for KIF2A and the indicated antibodies (scale bars: 0.5µm). Relative centriolar levels of KIF2A (normalized to CETN1, one-way ANOVA, \*\*\*\**P*<0.0001) are quantified in I. (J,K) hTERT RPE-1 TTBK2 KO cells expressing indicated Flag-TTBK2 constructs were transfected with control (NC) or KIF2A siRNA (scale bars: 2µm). The length of cilia (one-way ANOVA, \*\**P*<0.01, \*\*\**P*<0.001) is quantified in K.

**Fig 5:**
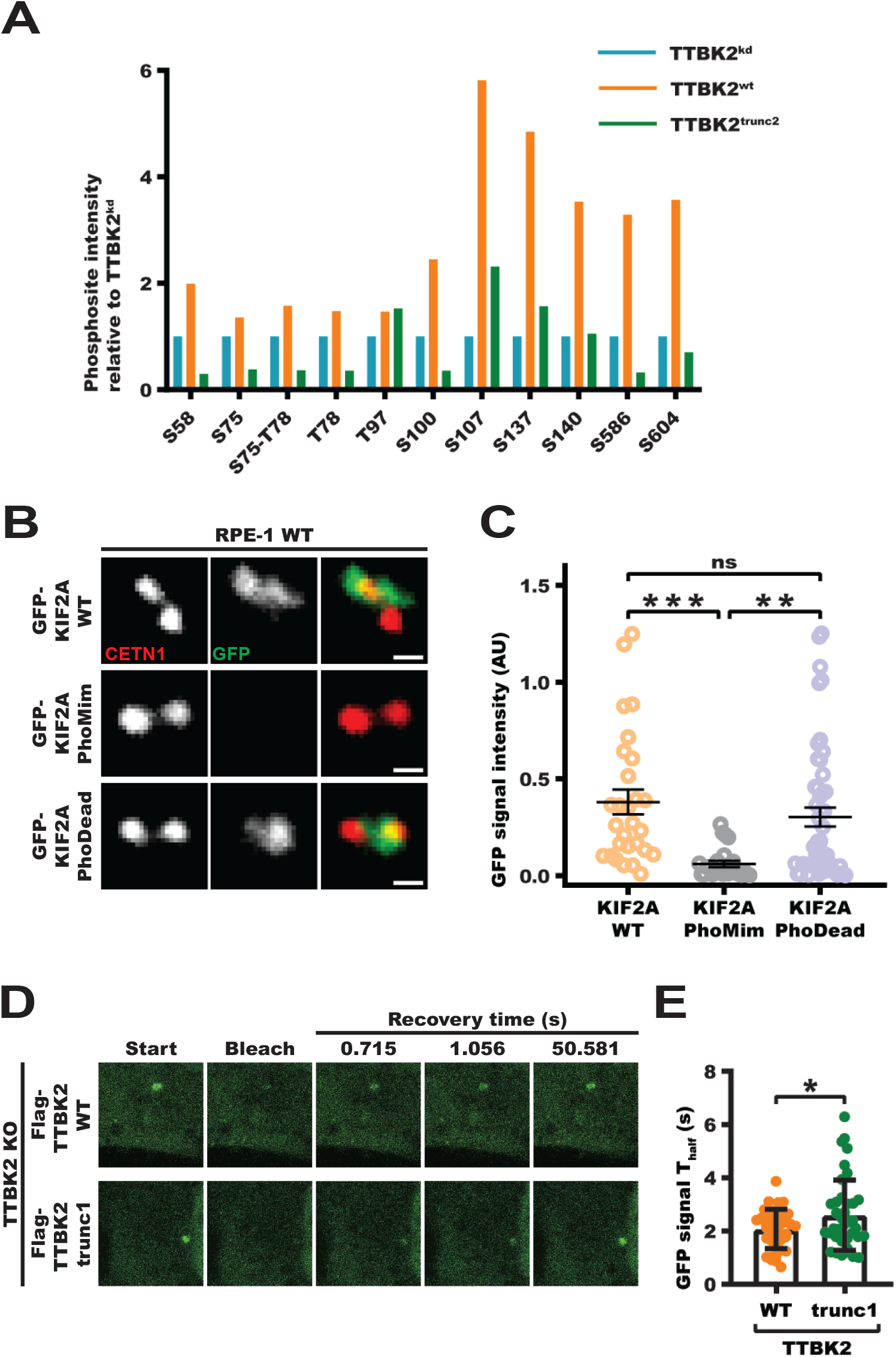
TTBK2-induced phosphorylations alter KIF2A basal body pool. (A) HEK293T cells were transfected with Flag-tagged KIF2A and GFP-tagged TTBK2 constructs, lysed, incubated with anti-Flag affinity beads, and the isolates were subjected to MS/MS analysis. Relative phosphorylation intensities were quantified as means of 3 independent experiments and normalized to TTBK2^kd^. (B,C) hTERT RPE-1 cells were transfected with GFP-KIF2A constructs, fixed, and stained for CETN1 (scale bars: 0.5µm). Relative centriolar GFP signal levels (normalized to CETN1, one-way ANOVA, \*\**P*<0.01, \*\*\**P*<0.001) are quantified in C. (D,E) hTERT RPE-1 cells expressing Flag-TTBK2 constructs were transfected with GFP-KIF2A^wt^ and subjected to FRAP analysis. Individual T _half_ (unpaired T-test, \**P*<0.05) are uantified in E.

**Figure 6:**
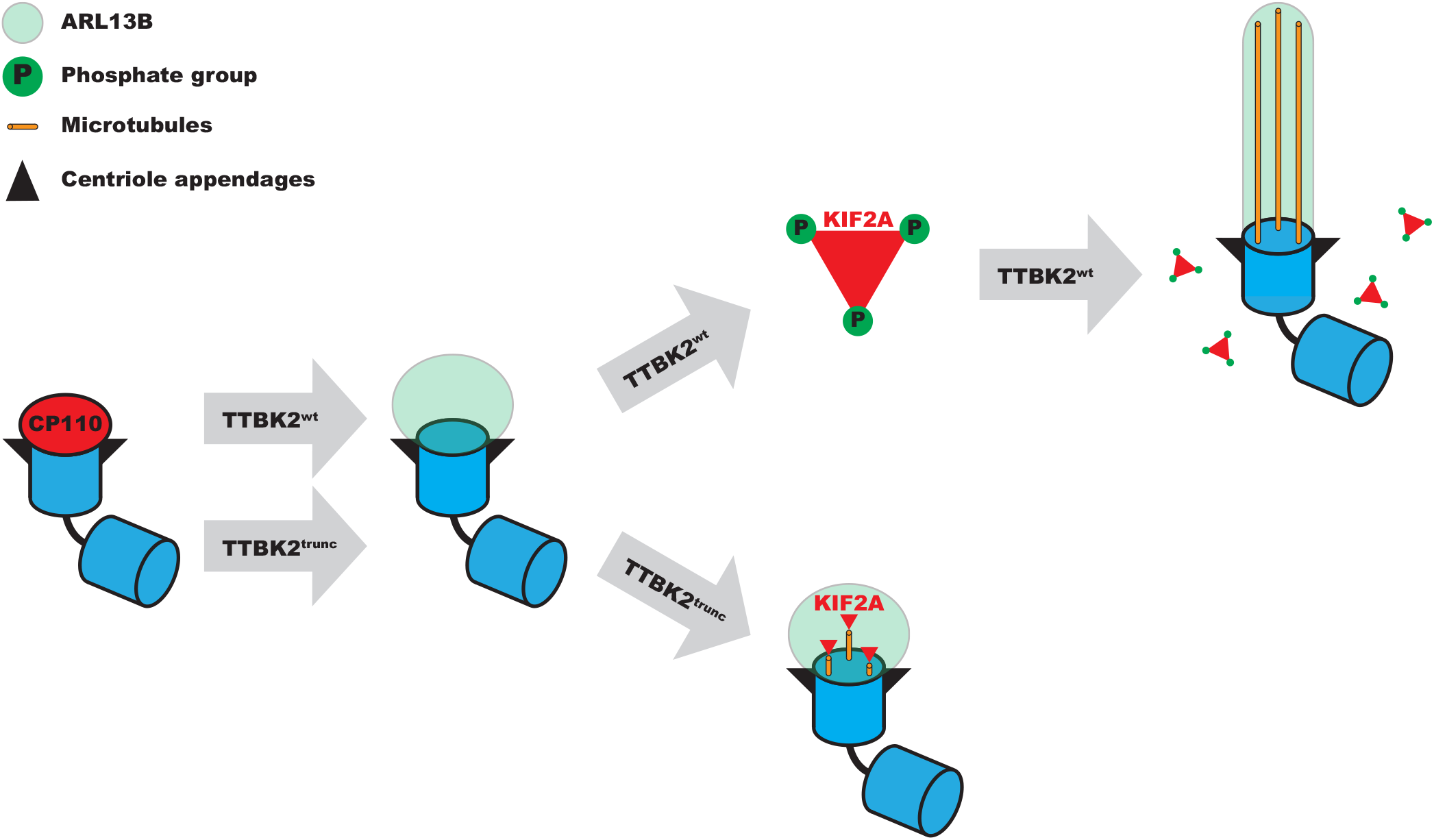
TTBK2 restrains KIF2A to promote cilia growth. In our model, TTBK2 kinase activity is required to prime the basal body for cilia outgrowth. Both TTBK2^wt^ and truncated TTBK2 constructs are sufficient to support this process. In addition, TTBK2^wt^ supports axoneme elongation by phosphorylating KIF2A at a specific cluster of serines, which reduces KIF2A basal body pool, inhibits KIF2A cilia-resorption activity, and ultimately promotes axoneme elongation. On the other hand, truncated TTBK2 constructs fail to inhibit KIF2A, which blocks proper axoneme growth.

## Discussion

TTBK2 has emerged as a crucial regulator of primary cilia formation, but the full scope of its activities and the underlying mechanisms are only starting to become apparent. Herein, we presented evidence that TTBK2 supports axoneme growth by restraining KIF2A levels at the MC.

Earlier work established that the MC-associated pool of KIF2A is phosphorylated by mitotic kinase PLK1 to trigger primary cilia resorption ^24^. Our data demonstrate that in addition to its MC-associated pool, KIF2A can also be detected along axonemal microtubules inside the cilium. This observation raises several intriguing possibilities. First, our expansion microscopy data suggest the MC-associated pool of KIF2A is well within the reach of TTBK2 kinase activity at MC^34,35^, given the flexibility of its long, highly unstructured C-terminal part^36^. Thus, KIF2A seems to be in a suitable position for direct regulation by TTBK2 phosphorylation, analogous to their interactions at microtubule plus ends^22^. Further, the presence of KIF2A inside the cilium helps to explain how KIF2A microtubule-depolymerization activity might mediate the resorption of primary cilia observed earlier^24^. However, we cannot exclude a contribution of additional mechanisms, e.g. KIF2A action towards microtubules anchored to the subdistal appendages of the MC^37,38^. We also noted that the axoneme-associated pool of KIF2A was somewhat easier to detect in the case of KIF2A^KVD^, perhaps due to altered dynamics of tubulin-KIF2A^KVD^ interaction^39,40^. Furthermore, our data suggest that altered KIF2A turnover contributes to primary cilia defects related to the activities of truncated TTBK2 moieties. We showed that KIF2A accumulated at the MC of TTBK2 KO hTERT-RPE-1 cells expressing TTBK2^trunc1/2^, and that primary cilia length can be rescued by KIF2A depletion. Noteworthy, similar accumulation of KIF2A has also been observed in TTBK2 hypomorphic mutant mice^19^.

We and others have demonstrated that the C-term region of TTBK2, missing in SCA11-related TTBK2 proteins, is essential for its interaction with CEP164^25,36,41^. Still, we were initially surprised to see the difference in TTBK2^trunc1/2^-induced phosphorylation between CEP164 and DVL3. We argue that such selectivity in targeting individual TTBK2 substrates may be explained by the hampered interaction capabilities of truncated TTBK2. Importantly, our rescue experiment using the CEP164 chimera demonstrates that the diminished ciliogenesis-promoting activity of TTBK2^trunc1/2^ resides in its spatial properties (miss-localization), rather than a defective kinase activity *per see*. Importantly, the expression of TTBK2^trunc1/2^ in hTERT RPE-1 TTBK2 KO allowed for semi-permissive conditions for cilia assembly. Intriguingly, we found that the initial events of primary cilia formation (e.g. formation of a membrane vesicle, CP110 removal, IFT recruitment), were rather moderately affected. In contrast, TTBK2^trunc1/2^ was not able to effectively mediate KIF2A turnover and support axoneme elongation. Based on these observations we conclude that individual steps of ciliogenesis likely differ in their requirements for TTBK2 activity. Furthermore, we speculate that concentrated TTBK2 activity at the MC is not strictly necessary for the phosphorylation of at least some of its basal body substrates, provided the levels and activity of the kinase outside of the MC are sufficiently high. In fact, our recent work has demonstrated that TTBK1, which does not localize to the MC, is able to partially support cilia formation in TTBK2 KO^42^, in line with the activities of TTBK2^trunc1/2^ reported here.

In our model (Figure 6), we conclude that the novel function of TTBK2 during cilia assembly and maintenance lies in destabilizing KIF2A basal body pool. In contrast, altered KIF2A turnover in the presence of truncated TTBK2 leads to KIF2A accumulation at the ciliary base, and in turn reduced cilia length. Furthermore, our data demonstrate that TTBK2 phosphorylation of the cluster S135-S140 in KIF2A contributes to that. Nonetheless, differential phosphorylation of additional substrates by TTBK2^wt^ vs truncated TTBK2 may also play a role in here. In sum, our data bring new insights into the biology of SCA11-related TTBK2 variants and the regulation of primary cilia formation.

The last (but not least) observation we find pertinent to discuss relates to TTBK2-mediated removal of CP110. As TTBK2^trunc1/2^ does not interact with CEP164, in principle it has no means to “distinguish” between MC- and DC-associated CP110. However, while elevated levels of TTBK2_trunc1/2_ over TTBK2_wt_ likely compensate for the absence of TTBK2_trunc1/2_ from MC, our data clearly show that TTBK2_trunc1/2_ can affect the MC-pool of CP110, but not the DC-pool of CP110. This “decoupling” of TTBK2 activity from the “recruiting” role of CEP164 challenges some of the recently proposed models of TTBK2-induced removal of CP110^43^ and suggests the existence of an additional mechanism ensuring MC-specific CP110 loss. We speculate that distal appendages may be plausible candidates for that function. Intriguingly, docking of vesicles to the distal appendages has been proposed to act upstream of CP110 removal^9,15,44–45^. However, other studies reporting defective vesicle docking have found no concomitant defect in the removal of CP110^46,47^. Thus, the cooperation of distal appendages and TTBK2 to regulate primary cilia assembly warrants further investigation.

## Materials and methods

### RPE-1 cell culture and stable line derivation

All hTERT RPE-1 cell lines were cultivated in DMEM/F12 cultivation media (Thermo Fisher Scientific, Cat.N.31331028), supplemented by 10% fetal bovine serum (Biosera), 1% penicillin/streptomycin and 1% L-glutamine. To induce primary cilia formation, the cells were cultivated in serum-free complete media for the last 24 hours of the experiment. Plasmid transfection was carried out using the Lipofectamine 3000 Transfection Reagent (Thermo Fisher Scientific, Cat.N. L3000001) and following the manufacturer’s manual. Paclitaxel treatment was carried out by adding paclitaxel (Merck, Cat.N. T7402) to a final concentration of 5µM and leaving it in the culture for 4 hours before fixing the cells.

To generate stable transgenic cell lines, hTERT RPE-1 Flp-In T-Rex cells (a gift from Erich A. Nigg) or hTERT RPE-1 Flp-In T-REX TTBK2 KO cells^16^ were seeded on 5cm dishes, grown to 90% confluency, and co-transfected with pOG44 (5μg total DNA) and a donor vector (pgLap1/2, 500ng total DNA) containing the gene of interest (GOI) coupled to a gentamycin-resistance gene. Transfectants with stably integrated GOI were then selected based on their resistance to G418 (0.5 mg/ml, 1-2 weeks, Merck, Cat.N. G8168). To induce the expression of the GOI the cells were treated with doxycycline (2 μg/ml, Merck, Cat.N. 3072) for the duration of the experiment. For a list of plasmids used in this work see Supplementary Table 1.

For KIF2A knockdown, the cells were seeded on glass coverslips in a 24-well format and cultivated in complete media. After 24 hours, the cells were transfected with KIF2A siRNA using Lipofectamine RNAiMAX (Thermo Fisher Scientific, Cat.N.13778100) and following the manufacturer’s manual. For siRNA details see Supplementary Table 1.

### HEK293T cell culture and transfection

HEK293T cells were cultivated in DMEM cultivation media (Thermo Fisher Scientific, Cat.N.31966047), supplemented by 10% fetal bovine serum (Biosera) and 1% penicillin/streptomycin. Transfection of HEK293T cells was carried out using polyethyleneimine (PEI) in the following way: PEI was incubated in serum-free DMEM media for 10 minutes, plasmids (Supplementary table 1) were equilibrated in serum-free DMEM media and then mixed with PEI in a 3μl of PEI to 1μg of plasmid ratio. The resulting mixes of plasmid and PEI in DMEM were then added to cells and left in the culture overnight, the media was changed for a fresh complete media on the next day.

### Western blot

Cells were lysed in 1x Laemli lysis buffer (62.5mM Tris-HCl pH 6.8, 2% 2-mercaptoethanol, 10% glycerol, 0.01% bromphenol blue, 2% sodium dodecyl sulfate (SDS)). SDS-PAGE and membrane transfer were performed using instrumentation by BioRad (Mini-PROTEAN tetra vertical electrophoresis cell, Mini Trans-blot module). Cell lysates were loaded to a 5% stacking gel combined with an 8% running gel and ran at 150V in running buffer (0.1% SDS, 0.192M glycine, 0.025M Tris-base). The proteins were transferred to an Immobilon-PVDF membrane (Merck, Cat.N. IEVH00005) at 100V for 75 minutes in transfer buffer (20% methanol, 0.192M glycine, 0.025M Tris-base). The membranes were then blocked in 5% solution of skimmed milk in wash buffer (20mM Tris-base, 0.1% Tween 20, 150mM NaCl) and incubated with primary antibodies (see Supplementary table 1) diluted in the same solution overnight at 4°C. The next day the membranes were washed 3 times for 10 minutes in wash buffer, incubated with secondary antibodies (Supplementary table 2) diluted in 5% milk/wash buffer for 2 hours at room temperature and then washed in wash buffer 3 times for 10 minutes. The membranes were then developed using ECL Prime (Merck, Cat.N.GERPN2236) and Chemidoc Imaging System (BioRad, Cat.N.12003154). For an easier analysis of membranes, a labelled protein ladder (Thermo Fisher Scientific, Cat.N.26625) was used in every SDS-PAGE performed.

### Immunoprecipitation and MS/MS analysis

To isolate KIF2A for the purposes of MS/MS analysis, HEK293 cells grown on 15cm plates were transfected with 2μg of Flag-KIF2A plasmid + 8μg of the corresponding GFP-TTBK2 construct. 48 hours post transfection the cells were scraped into lysis buffer (0.5% Triton X-100, 0.5% NP40, 150mM NaCl, 20mM Tric-HCl pH 7.4) containing Complete Mini Protease Inhibitor Cocktail (Roche, 1 tablet/10ml buffer) and lysed for 15 minutes on ice. The cell lysates were centrifuged at 16,000 x g for 10 minutes at 4°C and the supernatants were then incubated overnight at 4°C with anti-Flag M2 affinity gel beads (Merck, Cat.N. A2220). M2 beads were then pelleted and washed three times with lysis buffer containing protease inhibitors, resuspended in 1x Laemli lysis buffer and boiled at 95°C for 10 minutes. The resulting samples were separated on an 8% acrylamide gel, prominent bands of the correct size were cut out and subjected to protein extraction followed by MS/MS analysis^16^. The final plotted values represent mean relative phosphointensities from 3 independent experiments, only phosphosites detected with an absolute intensity above 10^6^ units were taken into consideration.

### Software and data analysis

All statistical analyses (one-way ANOVA with Tukey’s post-test or unpaired Student’s T-tests) were performed in GraphPad Prism software (version 8.0.1). The ACDC^48^ Matlab script (version 0.9) or the CiliaQ^49^ Fiji plugin (version 0.1.4) were used for measuring cilia number and length. 3D reconstructions of expansion microscopy images were generated using Imaris software (version 9.8.2).

### Immunofluorescence microscopy

hTERT RPE-1-derived cell lines or HEK293T cells were grown on glass coverslips in a 24-well format, briefly washed with PBS and fixed using ice-cold methanol at -20°C for 20 minutes. The coverslips were then briefly washed 2 times with PBS, incubated with primary antibodies overnight at 4°C, washed 3×10 minutes with PBS before being incubated with secondary antibodies for 2 hours in a dark chamber at room temperature, washed 3×10 minutes with PBS again and mounted using glycergel (Dako, Cat.N.C0563). The imaging was performed with the use of ZEISS microscopes, either with AxImager or LSM-800. For a list of antibodies used in this work see Supplementary Table 2.

Image analysis and signal intensity measurements were done in Fiji (version 2.0). Centriolar signal intensity was measured by drawing an ellipsoidal region of interest (ROI) around the centrioles and measuring mean signal intensities inside the ROI (= centriolar signal), then slightly moving the ROI next to the centrioles and measuring mean signal intensities again (= background signal). The final plotted values are equal to

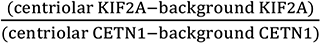

### Expansion microscopy

Following cultivation on glass slides in a 24-well format, the cells were briefly washed with PBS and fixed with fixation buffer (PBS, 4% parafolmaldehyde, 4% acrylamide) for 48 hours at room temperature, then briefly washed 2 times with PBS. For each sample, a droplet of polymerizing acrylamide gel (PBS, 19% sodium acrylate, 10% acrylamide, 0.1% N,N’-Methylenebisacrylamide, 0.5% ammonium persulfate, 0.5% temed) was prepared, the coverslips were quickly put on top of the gel droplet, with the cells facing the droplet. The samples were then incubated at 4°C for 10 minutes, then at 37°C for 30 minutes. The resulting coverslips covered in polymerized gel were transferred to a denaturation buffer (50mM Tris-base, 200mM NaCl, 200mM SDS) and the glass coverslips were gently removed from the gels using flat forceps. The gels were then incubated in denaturation buffer at 95°C for 2 hours and let to expand for 1 hour in ddH_2_O at room temperature. The gels were then cut into smaller pieces and incubated overnight at room temperature in primary antibodies (Supplementary Table 2) diluted 1:50 in blocking buffer (PBS, 2% BSA, 0.02% sodium azide). The next day the samples were washed 2×30 minutes with ddH_2_O and then incubated at room temperature overnight in secondary antibodies (Supplementary Table 2) diluted 1:500 in blocking buffer. The following day the samples were washed 2×30 minutes with ddH_2_O, placed in a glass-bottom microscopy dish (Ibidi) and analysed using the LSM-880 Airy2 microscope by ZEISS.

Centriole/axoneme lengths (3B) were measured by first reconstructing the AcTUB signal in 3D using the Surfaces function in Imaris, then using the Elipsoid Axis Length function in Imaris to calculate the length of 3D-reconstructed centrioles/axonemes.

### Live-cell imaging

Reporter hTERT-RPE-1 Flp-In T-Rex TTBK2 KO cell lines with DOX-inducible expression of Flag-TTBK2 constructs and constitutive mNeonGreen-ARL13B reporter expression were prepared as described in^50^. For the time-lapse live imaging experiment cells were seeded in DMEM/F12 medium, 10%FBS, 1%L-glutamine and Penicilin/streptomycin, supplemented with 1 μg/mL DOX on a 10-well glass-bottom CELLVIEW CELL CULTURE SLIDE, PS, 75/25 MM (Greiner Bio-One) at a high density (∼30.000 cells per well). 72h after seeding the medium was replaced for FluoroBrite DMEM with, 1%L-glutamine and Penicilin/Streptomycin supplemented with 1 μg/mL DOX to start the starvation of cells and induce cilia growth and then were immediately subjected to imaging. Slides with cells were equilibrated in the microscope environmental chamber for at least 30 minutes before imaging start. We used Elyra7 inverted microscope equipped with a super-resolution structured illumination microscopy (SIM) module with Plan-Apochromat 40x/1.4 Oil DIC M27. Z-stack images were taken every 15 minutes, resulting multi-scene .czi file was processed in Zen Black Software by SIM^2^ Method. The processed Z-stacks were projected in one layer by Maximal Orthogonal Projection in Zen Blue Software and individual Scenes were saved as .tiff files using Bio-Formats Importer plugin in Fiji. Cilia length was measured with the Segmented line tool in Fiji.

### FRAP

hTERT RPE-1 TTBK2 KO cells expressing Flag-tagged TTBK2 constructs were seeded in μSlide 8 Well High (Ibidi) chambers. The next day the cells were transfected with GFP-KIF2A^wt^, cultivated for another 24 hours, and then subjected to FRAP analysis using the LSM-880 microscope system by Zeiss – centriolar GFP signal was bleached with a strong laser pulse (100% laser intensity, 500ms) and the region of interest was imaged over the following 60s before measuring GFP signal intensity using the ZenBlack software. Thalf was calculated from individual FRAP recovery curves using the EasyFRAP web tool^51^.

## Acknowledgements

We thank Erich Nigg, Peter Jackson, Randall Moon, and Stephan Angers for sharing reagents with us. This work was supported by grant from Czech Science Foundation (22-13277S) to LC. DV was supported by grant MUNI/C/0026/2020 from Masaryk University. We acknowledge the core facility CELLIM supported by the Czech-BioImaging large RI project (LM2023050 funded by MEYS CR) for their support with obtaining scientific data presented in this paper. Furthermore, CEITEC Proteomics Core Facility is gratefully acknowledged for the obtaining of the scientific data presented in this paper. CIISB, Instruct-CZ Centre of Instruct-ERIC EU consortium, funded by MEYS CR infrastructure project LM2023042 and European Regional Development Fund-Project „UP CIISB”(No. CZ.02.1.01/0.0/0.0/18_046/0015974), is acknowledged for the financial support of the measurements at the CEITEC Proteomics Core Facility.

