## Supplementary Table 1 for "Tau-Tubulin Kinase 2 restrains microtubule-depolymerizer KIF2A to support primary cilia growth"

**Supplementary Table 1: Plasmids, primers, siRNA**

| TTBK2 construct | Aminoacids | Kinase activity | Basal body localization | Description |
|---|---|---|---|---|
| TTBK2$^{wt}$ | 1-1244 | yes | yes | full-length wild-type TTBK2 allele |
| TTBK2$^{kd}$ | 1-1244 | no | yes | full-length TTBK2, D136 mutated to A |
| TTBK2$^{trunc1}$ | 1-450 | yes | no | shortened canonical TTBK2 sequence, codons 1-450 |
| TTBK2$^{trunc2}$ | 1-450 | yes | no | 1-base insertion at nucleotide 1329, creating a STOP at codon 451 |
| CEP164 chimera | - | yes | yes | TTBK2$^{trunc1}$ fused to CEP164 aminoacids 1136-1460 |

| Plasmid | Source |
|---|---|
| pgLap2 Neo | Bernatik et al., 2020, N-terminal Flag tag |
| pgLap2 Neo TTBK2$^{wt}$ | Bernatik et al., 2020 |
| pgLap2 Neo TTBK2$^{kd}$ | Bernatik et al., 2020 |
| pgLap2 Neo TTBK2$^{trunc1}$ | cds cloned into pgLap2 NEO (Bernatik et al., 2020) |
| pgLap2 Neo TTBK2$^{trunc2}$ | adenosine insertion into pgLap2 Neo TTBK2$^{wt}$ at nucleotide 1329 using site-directed mutagenesis |
| pgLap2 Neo CEP164 chimera | cds cloned into pgLap2 NEO (Bernatik et al., 2020) |
| pgLap2 Neo KIF2A$^{wt}$ | cds cloned into pgLap2 NEO (Bernatik et al., 2020) |
| pgLap1 Neo | G418-resistance gene was cloned into pgLap1 (a gift from Peter Jackson lab, Addgene: 19702), N-terminal GFP tag |
| pgLap1 Neo TTBK2$^{wt}$ | Cajanek et al., 2014 |
| pgLap1 Neo TTBK2$^{kd}$ | Cajanek et al., 2014 |
| pgLap1 Neo TTBK2$^{trunc2}$ | adenosine insertion into pgLap2 Neo TTBK2$^{wt}$ at nucleotide 1329 using site-directed mutagenesis |
| pgLap1 Neo KIF2A$^{wt}$ | cds cloned into pgLap1 NEO |
| pgLap1 Neo KIF2A$^{KVD}$ | KVD domain mutated to triple A using site-directed mutagenesis |
| pgLap1 Neo KIF2A$^{PhoMim}$ | S135, S137, and S140 mutated to G using site-directed mutagenesis |
| pgLap1 Neo KIF2A$^{PhoDead}$ | S135, S137, and S140 mutated to A using site-directed mutagenesis |
| DVL3 | a gift from S. Angers (doi: 10.1038/ncb1381) |
| Myc-CEP164 | Cajanek et al., 2014 |
| Topflash | Addgene: 12456 |
| Renilla | Addgene: E2241 |
| BCAT | Orford et al., 1997 |

| Primer | Sequence | Description |
|---|---|---|
| TTBK2_trunc2_fwd | ttccactggtgcgaaagttaacgttccattcacag | mutagenesis primer for adenosine insertion at nucleotide 1329 |
| TTBK2_trunc2_rev | ctgtgaatggaacgttaactttcgcaccagtggaa | mutagenesis primer for adenosine insertion at nucleotide 1329 |
| KIF2A_KVDmut_fwd | aaatgtttggttttctaggtaccttgttaaagctgctgcttgttttggttcatgtaccatcacaacatct | mutagenesis primer for mutating KIF2A KVD domain to triple A |
| KIF2A_KVDmut_rev | agatgttgtgatggtacatgaaccaaaacaagcagcagctttaacaaggtacctagaaaaccaaacattt | mutagenesis primer for mutating KIF2A KVD domain to triple A |
| KIF2A_PhoMim_fwd | ggggtccaaattcctttttgcagcttgaactggctctatatcctcaacctcaccattctgttgtgcagaggaagactgttcaggaaat | mutagenesis primer for mutating S135, S137, and S140 to G |
| KIF2A_PhoMim_rev | atttcctgaacagtcttcctctgcacaacagaatggtgaggttgaggatatagagccagttcaagctgcaaaaaggaatttggacccc | mutagenesis primer for mutating S135, S137, and S140 to G |
| KIF2A_PhoDead_fwd | ttttgcagcttgaactggagctatatctgcaacagcaccattctgttgtgcagagg | mutagenesis primer for mutating S135, S137, and S140 to A |
| KIF2A_PhoDead_rev | cctctgcacaacagaatggtgctgttgcagatatagctccagttcaagctgcaaaa | mutagenesis primer for mutating S135, S137, and S140 to A |

| siRNA | Description |
|---|---|
| siKIF2A | Thermo Fisher Scientific, Cat.N. 4390824 |
| siNC | sense: 5' CGUACGCGGAAUACUUCGATT 3', antisense: 3' TTGCAUGCGCCUUAUGAAGCU 5' |
