## Supplementary Table 2 for "Tau-Tubulin Kinase 2 restrains microtubule-depolymerizer KIF2A to support primary cilia growth"

**Supplementary Table 2: Antibodies**

| Antibody | Manufacturer/Cat.N. | Working concentration |
|---|---|---|
| AcTUB | Merck/00020913 (C3B9) | 1:50 (Expansion microscopy) |
| ARL13B | Proteintech/17711-1-AP | 1:1000 (IF), 1:50 (Exp microscopy) |
| Centrin1 | Proteintech/12794-1-AP | 1:50 (directly labelled with Alexa647) |
| CAP350 | Thermo Fisher/A301-170A | 1:50 (directly labelled with Alexa647) |
| CEP135 | Proteintech/24428-1-ap | 1:500 |
| CP110 | Proteintech/12780-1-ap | 1:500 |
| Flag | Merck/F1804 | 1:500 |
| α-TUB | Proteintech/66031-1-ig | 1:2000 |
| γ-TUB | Merck/T6557 | 1:500 |
| IFT88 | Proteintech/13967-1-AP | 1:500 |
| KIF2A | Santa Cruz/sc271471 | 1:250 |
| TTBK2 | Merck/HPA018113 | 1:1000 |
| GFP | Proteintech/50430-2-ap | 1:50 (Exp microscopy) |
| DVL3 | Proteintech/13444-1-AP | 1:500 |
| Alexa 488 anti-mouse | Thermo Fisher/A-21202 | 1:1000 |
| Alexa 488 anti-rabbit | Thermo Fisher/A-21206 | 1:1000 |
| Alexa 568 anti-mouse | Thermo Fisher/A10037 | 1:1000 |
| Alexa 568 anti-rabbit | Thermo Fisher/A10042 | 1:1000 |
| Alexa 647 labelling kit | Thermofisher/A20186 | manufacturer's manual |
| HRP-linked anti-rabbit IgG | Cell Signalling/7074S | 1:2000 (WB) |
| Peroxidase-linked anti-mouse IgG | Sigma/A4416 | 1:2000 (WB) |
