## Supplementary Figures for "Tau-Tubulin Kinase 2 restrains microtubule-depolymerizer KIF2A to support primary cilia growth"

### Supplementary Figure 1

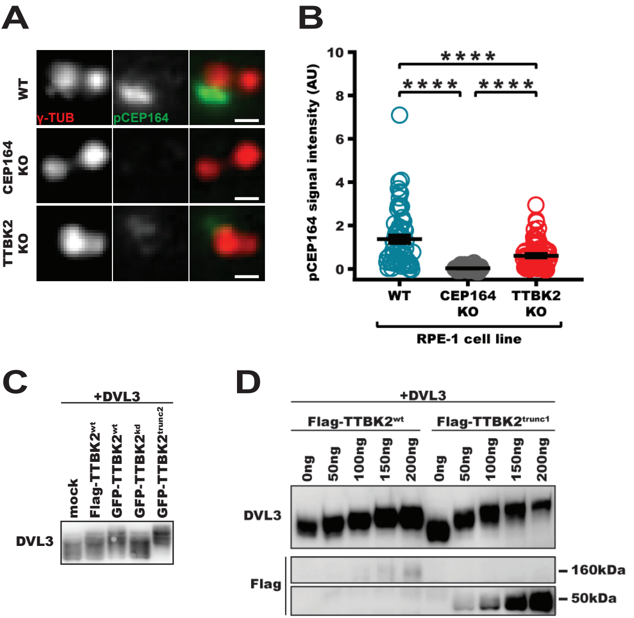


**Suppl. Fig. 1: Validation of the pCEP164 antibody and the activity of TTBK2$^{trunc2}$**

(A,B) Wild-type, CEP164 KO, and TTBK2 KO hTERT RPE-1 cells were fixed and stained for pCEP164 (scale bars: 0.5μm.) Relative centriolar levels of pCEP164 (normalized to γ-TUB, one-way ANOVA, ****$P$<0.0001) are quantified in B. (C,D) HEK293T cells were co-transfected with DVL3 and TTBK2 constructs, lysed, and subjected to WB analysis. (C) Similar to Flag-TTBK2$^{trunc1}$, GFP-TTBK2$^{trunc2}$ induced a significant mobility shift in DVL3. (D) Dose-dependent effect of TTBK2 variants on DVL3. Note that TTBK2$^{wt}$ and TTBK2$^{trunc1}$ show comparable effect when expressed at similar levels (200ng vs 50ng respectively).

### Supplementary Figure 2

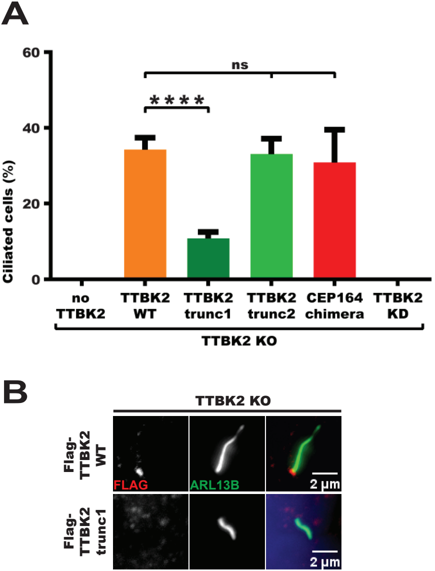


**Suppl. Fig. 2: Ciliogenesis rescue by TTBK2 constructs**
(A) Cilia number (one-way ANOVA, ****$P$<0.0001) was determined in hTERT RPE-1 cells expressing indicated Flag-TTBK2 constructs. (B) HEK293T TTBK2-KO cells were transfected with Flag-TTBK2 constructs, fixed, and stained for ARL13B and the indicated antibodies. Both Flag-TTBK2$^{wt}$ and Flag-TTBK2$^{trunc1}$ rescued cilia formation.

### Supplementary Figure 3

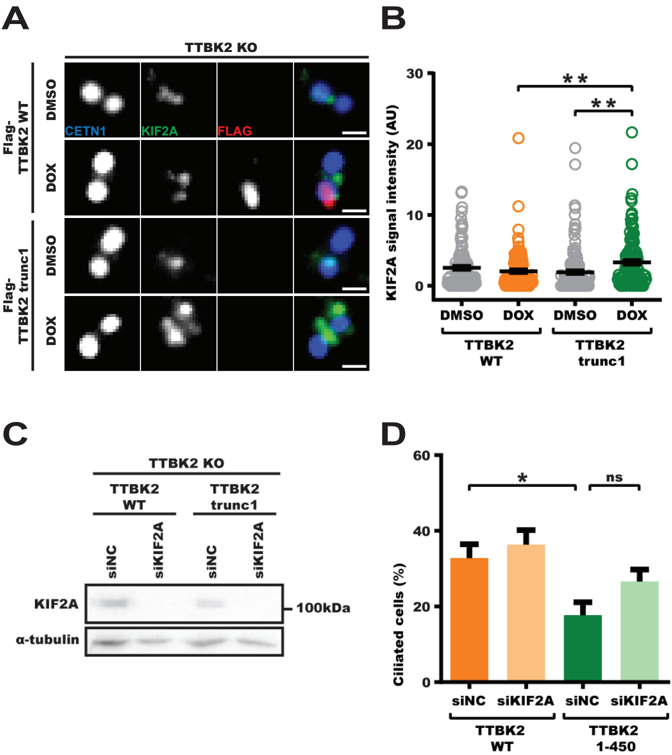


**Suppl. Fig. 3: KIF2A accumulation and KIF2A knockdown**

**(**A,B) hTERT RPE-1 cells expressing Flag-TTBK2 constructs were treated either with DMSO (control group) or DOX, fixed, and stained for KIF2A and the indicated antibodies. Relative levels of KIF2A (normalized to CETN1, one-way ANOVA, ***P*<0.01) were quantified in B. (C) Validation of KIF2A knockdown efficacy. hTERT RPE-1 cells expressing Flag-TTBK2 constructs were transfected either with mock (siNC) or KIF2A (siKIF2A) siRNA. After 48 hours, the cells were lysed and subjected to WB analysis. (D) The percentage of ciliated cells (one-way ANOVA, **P*<0.05) was determined in hTERT RPE-1 TTBK2 KO cells expressing indicated Flag-TTBK2 constructs following siRNA transfection.
